## Supplementary Data for "qByte: Open-source isothermal fluorimeter for democratizing analysis of nucleic acids, proteins and cells"

### Supplementary Data 1

#### qByte: Open-source isothermal fluorimeter for democratizing analysis of nucleic acids, proteins and cells

Francisco J. Quero, Guy Aidelberg, Hortense Vielfaure, Yann Huon de Kermadec, Severine Cazaux, Amir Pandi, Ana Pascual-Garrigos, Anibal Arce, Samuel Sakyi, Urs Gaudenz, Fernan Federici, Jennifer C. Molloy, Ariel Lindner

October 28, 2024

#### Summary

This supplementary information provides additional data on: (1) hardware details, including insights obtained during the prototyping process, the Bill of Materials (BoM) for the components used, and the testing and calibration data for the thermal and optical modules; (2) a detailed description of the software's functionality and structure, covering both the embedded system and the user interface; and (3) the sequences of all DNA fragments used in the experiments.

For the most detailed and up-to-date instructions on replicating the system, readers are directed to the project's GitLab repository. The repository also includes all the necessary manufacturing files, such as Gerber files for electronic boards and 3D printing files.

#### Links

1. Repository
   (a) BoM
   (b) 3D printing files
   (c) Gerber files
   (d) Embedded system source
2. OnShape 3D models
3. EasyEDA project

### Hardware

#### S1A Data - Prototyping insights

The qByte was developed with a modular iterative design process, where we built and tested each subsystem step by step. This approach helped us improve its design and simplify its production: we report here the lessons learned by building our system as we believe they can be more generally useful in the design and production of other devices. This section briefly covers the key insights from our process, with more technical details available in the previously linked Gitlab repository.

**Heated lid:** In the early stages of prototyping, the lack of a heated lid caused reagent evaporation, increasing the reagents concentration (primers and dies) and generating a saturated background signal. To solve this, we initially used a layer of mineral oil as a seal, which effectively reduced evaporation and allowed us to achieve our first results. However, this method was not applicable for experiments that required gas exchange, such as those using lysate or cell-free samples where oxygen exchange is critical. This made us finally opt for a heated lid implementation as a more general solution to prevent evaporation.

Please note that a layer of mineral oil should still be added in experiments where reaction temperature was set at high temperatures ($> 70$°C), as then the heated lid needs to reach a temperature of around 120°C. In this scenario there is a much greater gradient between the heaters (lid and tube holder) and the room temperature than between the lid and the tube holder, so the lid heating pressure becomes inefficient. Ultimately this makes the central area of the tube (far from both the tube holder and the lid) approach room temperature and become colder than the tube holder itself, therefore generating a ring of condensation in that region.

We included the lid as a separate flexible PCB with an aluminum stiffener, allowing us to integrate the heating resistors, the sensor and the connector into a single component.

**Temperature measurement:** We developed several versions of the tube holder and heating PCB before achieving optimal thermal uniformity. Ultimately we used multiple heating segments to compensate for heat loss, and we included decoupling capacitors near the analog reading paths to filter out noise introduced by the power lane switch.

**Optical signals:** We tested different architectures for the analog measurement circuit. We first experimented with solutions involving an operational amplifier for each photodiode, minimizing their distance to the sensors, as a longer path length potentially allowed for the entry of electromagnetic noise (from, for example, the first coil-based heating lid designs). However, we realized that the internal component variability among the lower priced operational amplifiers was much greater than any electromagnetic noise induced, leading us to centralize all the measurements in a single, more reliable operational amplifier. To do so, we employed low resistance multiplexers to switch the different signals over the operational amplifier. To find the optimal sensing unit, we characterized multiple combinations of photodiodes and filters with a range of different concentrations of fluorescein and different controls (Fig. 3). In the mechanical part, we realized that if there is an empty space below, the PCB bends downwards, causing the photodiodes on the bent part to be slightly further away, reducing the signal they receive (as it decreases with the square of the distance from the light source). We optimized the size of the light-entry and fluorescence-exit holes, finding that the amount of LEDs light received by the reactions is critical, and determining the measured fluorescence intensity in a nonlinear way, and making the error difficult to calibrate by software.

**Component availability and quality:** We conducted continuous assessment of component availability, cost, and quality variations to inform decisions related to batch production. This was particularly crucial during the COVID-19 pandemic chip scarcity, which ultimately made us learn which components are more prone to shortage during a crisis.

**Optimizing case production for 3D printing:** The piece composed by the main body and cap, is specially designed to be printed as a single unit, including an internal hinge that allows movement between the cap and body, eliminating the need for any additional, non-printed components.

#### S1B Data - Components

The complete BoM, including descriptions, prices, and suppliers, can be found in the project's GitLab repository listed above. Fig. 1 presents a general price breakdown of the different components and their respective contributions to the total cost.

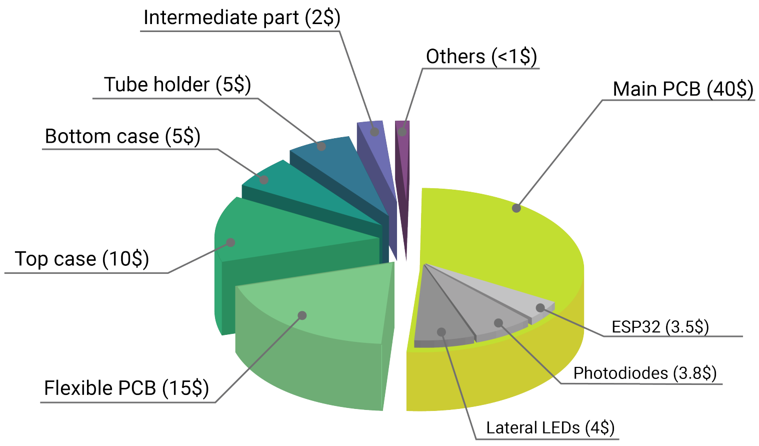


Figure 1: qByte component cost by category.

#### S1C Data - Calibration

##### Temperature

We performed the initial temperature calibration using an external thermometer with a thermocouple (PRO RS41, RS components), immersed in a tube filled with mineral oil. The system was calibrated first with the lid turned off to prevent heat transfer from the lid to the thermocouple wire at the top of the tube, which could have altered the temperature distribution along the wire and compromised the accuracy of the measurement.

After completing the initial calibration, we evaluated temperature variability across the eight samples by performing a melting curve analysis using a 17-nucleotide double-stranded DNA probe (predicted melting temperature of 55°C). The results were compared with those obtained from a commercial thermocycler (BioRad CFX96). We observed a temperature variability of approximately 1°C in the open qByte, compared to a 2°C variability in the commercial thermocycler (Fig. 2).

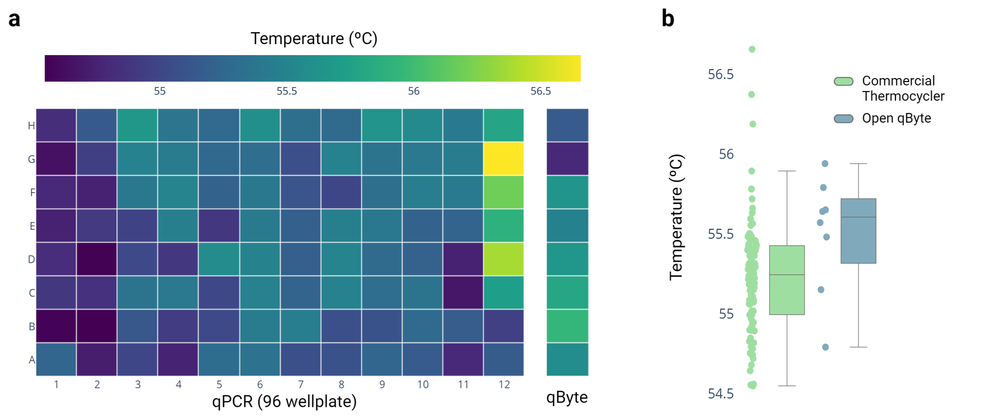


Figure 2: Temperature calibration results, including a heatmap of the melting peaks obtained for the melting probes using a commercial thermocycler and the qByte (a), as well as a boxplot of the results (b).

Once the temperature model values were adjusted for the first device, further recalibration with the external sensor did not lead to significant performance improvements in subsequent devices (data not shown). Consequently, as of this study, we have produced more than 20 devices that perform consistently well without the need for recalibration. For each device, we retained the initial calibration parameters and verified performance by conducting a melting curve test using the same melting probes.

Additionally, for applications requiring more stringent temperature calibration, we believe that DNA melting probes could serve as an efficient method to calibrate the devices. By incorporating different probes into a single tube and using it as an auto-calibration sample, the device could incrementally increase the temperature until detecting the melting peaks of the probes. It could then correct small discrepancies between devices by recalibrating the thermistor's resistance-to-temperature function based on the Steinhart-Hart model.

#### Fluorescence

We conducted two tests to characterize the fluorescence measurement system:

- **Photodiode Comparison**: We compared the performance of four SMD photodiodes, selected based on compatibility with the qByte design principles in terms of size and price. Each photodiode was tested in separate cells, measuring the signals received under two different filters—one orange and one green—resulting in one measurement per photodiode and filter.
- **Variability Characterization**: Using the two best-performing photodiodes, we used two devices (16 photodiodes in total) to characterize the replicability of the dynamic range across the 8 samples. This was performed using different concentrations of fluorescein (FAM), positive and negative controls of LAMP reactions, and samples of GFP production in cell lysate (Fig. 3).

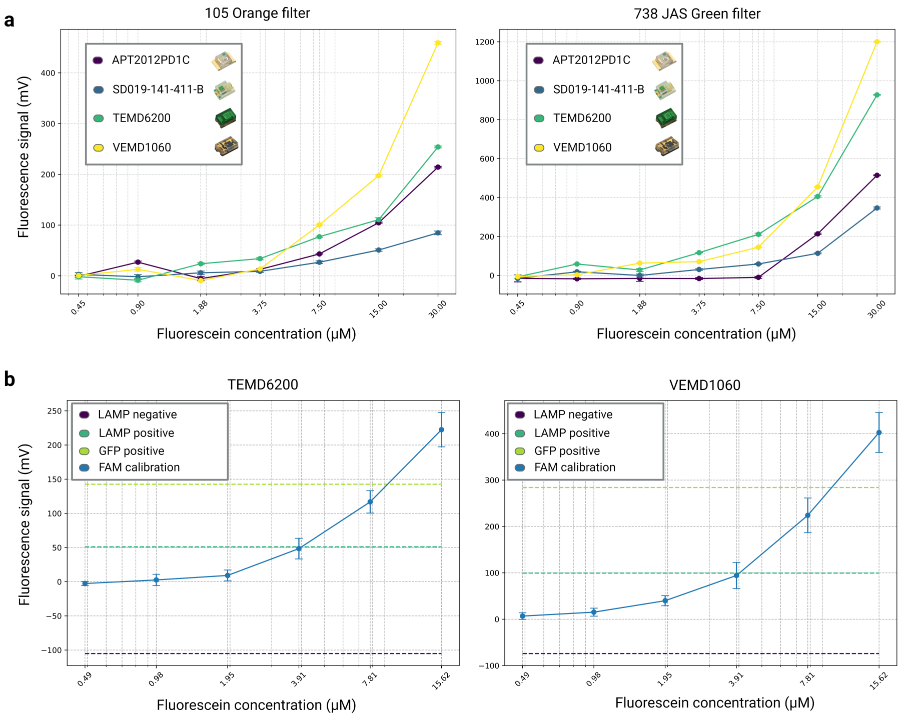


Figure 3: Fluorescence detection range results, including a test of different photodiode and filter combinations in a test cell (a), and a test with 16 replicates across two devices using the two photodiodes with the best signal, characterized by a FAM concentration curve, LAMP reaction controls, and GFP production in cell lysate (b).

### Software

The qByte software is divided into two components: the embedded system, which controls the hardware's operational logic, and the user interface, which is stored on the device's Serial Peripheral Interface Flash File System (SPIFFS)-mounted memory. The user interface is accessed as a web page through the user's browser when they connect to the device via the local network.

The code, binaries, and a guide for uploading them to the device can be found in the project's GitLab repository (see "Links" section).

#### S1D Data - Embedded System

The embedded system operates using an ESP32 microcontroller programmed in the Arduino language. Fig. 4 depicts the general logic of the system. The code is structured around the main ‘Setup()‘ and ‘Loop()‘ functions, with multiple asynchronous operations for command management, sensor reading, and temperature control.

During the ‘Setup()‘ phase, the system initializes key components:

- The serial port is configured to allow communication.

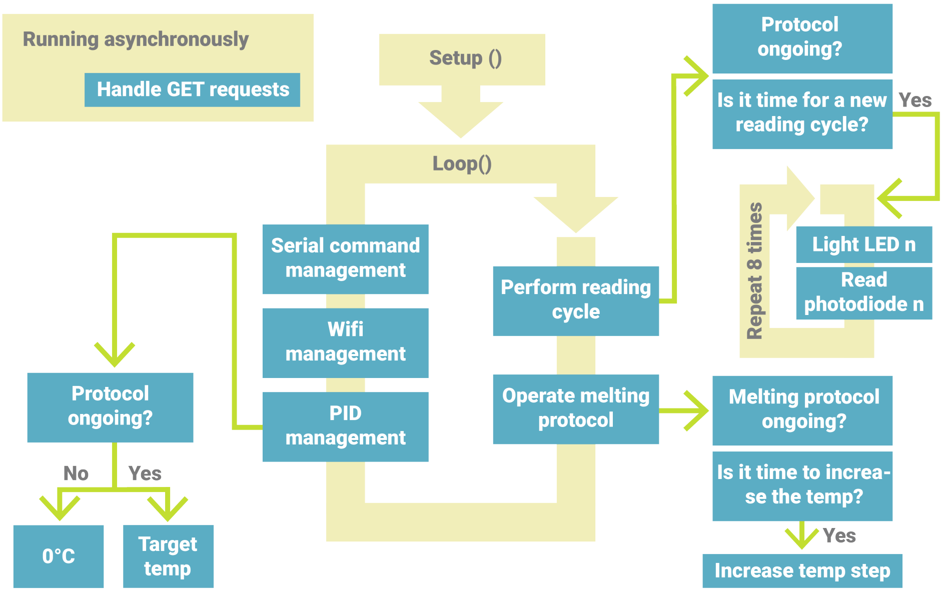


Figure 4: Diagram of qByte's simplified code structure.

- The SPIFFS file system is mounted to handle local storage, where the User Interface (UI), configuration, and results are stored.
- WiFi and Multicast DNS (mDNS) services are initialized, allowing the device to be accessed via a fixed local URL (qByte.local by default).
- A PID (Proportional-Integral-Derivative) control system is set up for accurate temperature regulation during experimental protocols.

The 'Loop()' function controls the continuous operation of the system. Different functions check during the loop whether it's time to operate, and if so, they execute and update their state. The management of GET requests is handled by interrupts using the asynchronous library "AsyncTCP". The core logic during a protocol revolves around checking the status of the experimental protocol, adjusting the temperature, and performing sensor readings when necessary.

- **Serial, WiFi**: Every cycle, qByte checks if there are new Serial commands to process and whether the WiFi is connected. If WiFi is disconnected, it attempts to reconnect every 60 seconds. The WiFi credentials are stored in the 'credentials.txt' file within the SPIFFS system and are reconfigurable via serial commands.
- **PID management**: Every cycle, if a protocol is running, qByte reads the current temperature of the lid and heating segments. The PID control system updates the PWM digital pins to smoothly approach the target temperature.
- **Protocol status check**: The system checks whether an experimental protocol is ongoing. If no protocol is active, qByte sets the temperature to 0°C. If a protocol is active, the PID updates the output of a digital PWM pin to smoothly approach the target temperature.

- **Reading cycle**: If a protocol is active and it is time for a new reading cycle, the system performs a sequence of sensor readings. This includes sequentially lighting LEDs and reading the corresponding photodiode signals, collecting data from the samples.
- **Melting protocol management**: During the melting protocol, the system monitors whether it's time to move to a new step and increase the temperature. If it is, the temperature is increased by a user-defined step size.

#### S1E Data - User interface

The UI is a web application programmed in HTML5 and JavaScript, stored on a file system mounted within the microcontroller (in the SPIFFS-mounted memory), and served over the local network as a web page. Once accessed, the web application runs on the user's mobile device, laptop, or desktop computer. To retrieve real-time data from the device, the UI makes HTTP GET requests to specific addresses, which return the device status or sample data in JSON format. The main structure of the UI is shown in Fig. 5 and described in more detail in the GitLab repository.

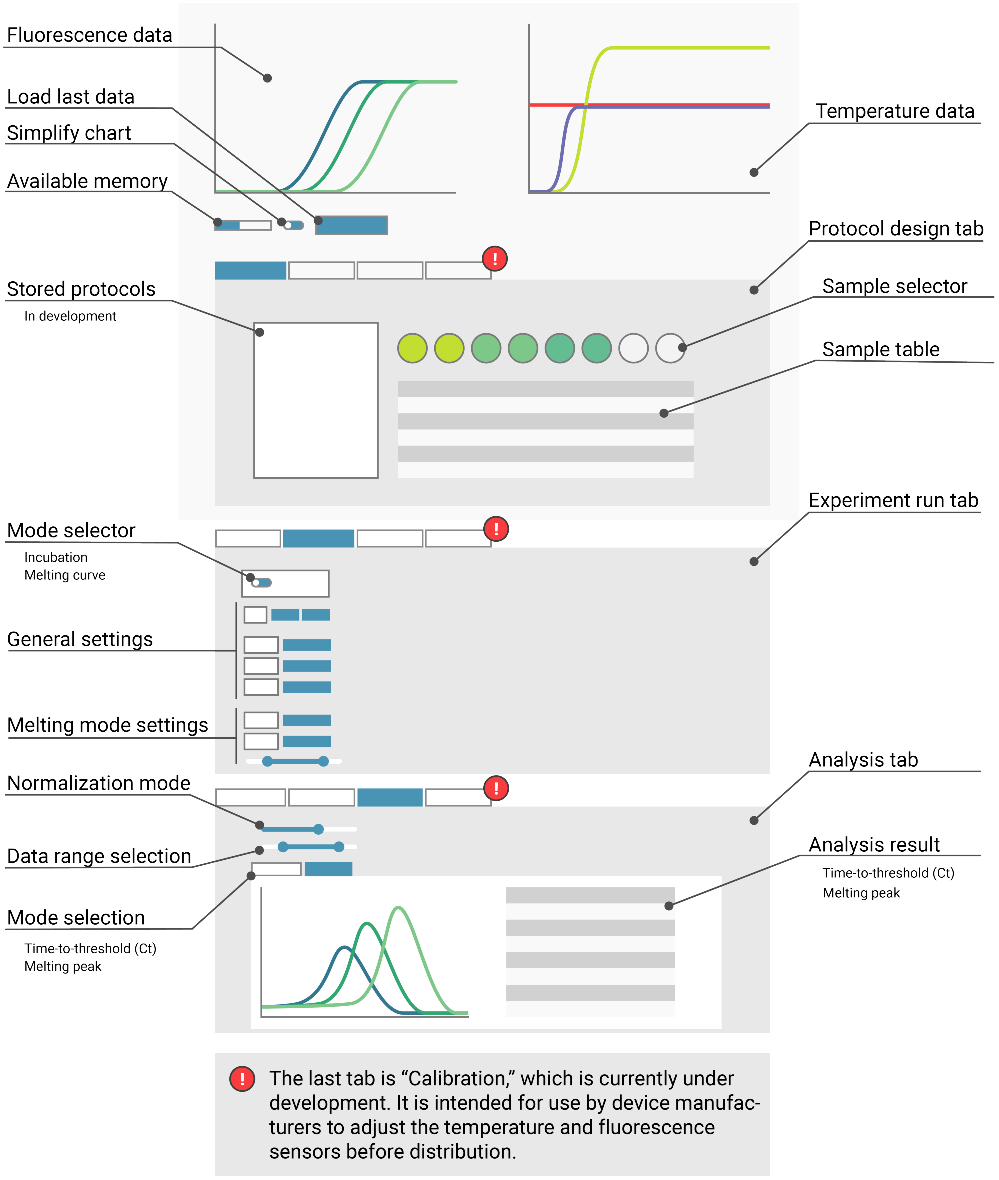


Figure 5: Overview of the qByte UI.

### 1 S1G Data - Sequences

| Name | Sequence | Type | Reference |
|---|---|---|---|
| **LAMP Primers** | | | |
| GMO detective | | | [1] |
| Coronadetective | | | [2] |
| *S. mansoni* | | | [3] |
| **CRISPR Tests** | | | |
| Sty16_B gRNA | UAAUUUCUACUCUUGUAGAUAAGGCAA<br>GUCUAAAUAUUG | RNA | |
| Target | CCCGGTCAAACTTAAGTTCCCAATATT<br>TAGACTTGCCTTTAAAAGATACCAGAG<br>CCCGA | DS | |
| Reporter | /56-FAM/TTATT/3IABkFQ/ | DS | |
| **Toehold-mediated translation** | | | |
| Toehold switch and trigger | | | [4] |
| **Melting probes (Size based)** | | | |
| 90bp, Tm:73 ºC | GTATGGTACTGAAGATGATTACCAAGG<br>TAAACCTTTGGAATTTGGAGCCACTTC<br>TGCTGCTCTTCAACCTGAAGAAGAGCA<br>AGAAGAAGA | DS | |
| 46bp, Tm:65 ºC | GTATGGTACTGAAGATGATTACCAAGG<br>TAAACCTTTGGAATTTGGAC | SS | |
| 26bp, Tm:54 ºC | GTATGGTACTGAAGATGATTACCAAG | DS | |
| 19bp, Tm:43 ºC | GTATGGTACTGAAGATG | DS | |
| 13bp, Tm:33 ºC | GTATGGTACTGAA | DS | |
| **Melting probes (Mutation based)** | | | |
| 46bp, reverse, 1 mismatch | GTCCAAATTCCAAAGGTTTACCTTAGT<br>AATCATCTTCAGTACCATAC | SS | |
| 46bp, reverse, 2 mismatches | GTCCAAATTCCAAAGGTTTACCTTAGT<br>AATCATCTTCAATACCATAC | SS | |
| 46bp, reverse, 3 mismatches | GTCCAAATTCCAAAGATTTACCTTAGT<br>AATCATCTTCAATACCATAC | SS | |
| 46bp, reverse, 0 mismatches | GTCCAAATTCCAAAGGTTTACCTTGGT<br>AATCATCTTCAGTACCATAC | SS | |
| **RBS (Application 3)** | | | |
| *In silico* generated | ACTGTTTTGTTAAACTTAAGGATAATT<br>TGT | DS | |
| B0033 & B0035 | | DS | [5] |

Table 1: **DS:** Double stranded. **SS:** Single stranded. **RNA:** Transcribed RNA from DNA template.

### 2 Supplementary references

#### References

[1] Guy Aidelberg. *Towards Democratization of Nucleic Acid Detection.* These de doctorat, Université Paris Cité, March 2021.

[2] Guy Aidelberg, Rachel Aronoff, Tatiana Eliseeva, Francisco Javier Quero, Hortense Vielfaure, Martin Codyre, Kathrin Hadasch, and Ariel B. Lindner. Corona Detective: A simple, scalable, and robust SARS-CoV-2 detection method based on reverse transcription loop-mediated isothermal amplification. *Journal of biomolecular techniques: JBT*, 32(3):89–97, September 2021.

[3] Isaac Owusu-Frimpong, Linda B. Debrah, samuel.k.armoo, Edward J. Tettevi, Yvonne A. Ashong, Naa A. Kuma, Frank Twum Aboagye, Bright K. Idun, and Mike Y. Osei-Atweneboana. Schistosome DNA Isolation and Colorimetric LAMP protocol for the detection of Schistosomiasis. December 2022.

[4] Arce Medina and Aníbal Andrés. Open platform for the implementation of RNA sensing reactions in cell-free systems. 2021.

[5] Ribosome Binding Sites/Prokaryotic/Constitutive/Community Collection - parts.igem.org. https://parts.igem.org/Ribosome_Binding_Sites/Prokaryotic/Constitutive/Community_Collection.
